# Graphmap2 - splice-aware RNA-seq mapper for long reads

**DOI:** 10.1101/720458

**Authors:** Josip Marić, Ivan Sović, Krešimir Križanović, Niranjan Nagarajan, Mile Šikić

## Abstract

In this paper we present Graphmap2, a splice-aware mapper built on our previously developed DNA mapper Graphmap. Graphmap2 is tailored for long reads produced by Pacific Biosciences and Oxford Nanopore devices. It uses several newly developed algorithms which enable higher precision and recall of correctly detected transcripts and exon boundaries. We compared its performance with the state-of-the-art tools Minimap2 and Gmap. On both simulated and real datasets Graphmap2 achieves higher mappability and more correctly recognized exons and their ends. In addition we present an analysis of potential of splice aware mappers and long reads for the identification of previously unknown isoforms and even genes. The Graphmap2 tool is publicly available at https://github.com/lbcb-sci/graphmap2.

## I. INTRODUCTION

The advances in sequencing technology, achieved by companies such as Oxford Nanopore technologies (ONT) [1] and Pacific Biosciences (PacBio) [2] resulted in production of long reads with over 10 kbp in length. Initially, such long reads had very high error rate which has steadily declined and the last generation of PacBio devices produce reads comparable in accuracy to Illumina short reads [3]. In addition, ONT has recently announced new R10 pores and their, in house, results are promising. Although in the field of RNA-seq short reads are still predominantly used, there is a need for longer reads which help in detection and quantification of isoforms. Algorithmically, mapping RNA-seq reads to known transcripts is equivalent to mapping DNA reads. Yet, mapping these reads to eukaryotic genomes is more complex due to RNA splicing. There are several RNA-seq splice-aware mapping tools developed for long reads produced by third generation sequencing technologies. An evaluation of these tools was given in [4] and [5], with GMAP [6] and Minimap2 [7] being the best performing tools. However, there is still room for improvement, especially in correctly aligning exon edges and finding all exons in transcripts - especially shorter exons.

In this work we present Graphmap2, an extended version of the Graphmap tool. Graphmap is a highly sensitive tool for mapping DNA reads to a reference genome, and we have extended it with the ability to map RNA reads. This extended version uses the same five-stage ‘read-funneling’ approach as the initial version [8] and adds upgrades specific for mapping RNA reads. The performance and accuracy of Graphmap2 was evaluated on simulated and real datasets and the results were compared to the state-of-the-art RNA mapping tools. In addition we analysed identification of potentially new isoforms and genes using long reads.

## II. METHODS

Figure 1 shows the complete process of aligning RNA reads to a reference genome. We can divide the original Graphmap mapping method into two stages; the first stage includes finding candidate positions on the reference genome using short seeds, and the second stage calculates the exact alignment of reads using Edlib - a fast implementation of Myers’ bit-vector algorithm [9]. The first stage which includes steps: (i) *region selection*, (ii) *graph mapping* and (iii) *longest common subsequence in k length substrings (LCSk)* [10] is kept in Graphmap2. For every read in the input dataset, this stage produces a set of approximate matches between parts of the read and parts of the reference. These matches are represented by anchors, where every anchor consists of start and end locations on the read, and start and end locations on the reference in that match. Graphmap2 then proceeds by extending the workflow with the following stages: (iv) anchor filtering (knapsack), (v) anchor alignment, (vi) alignment tuning and (vii) exon grouping and adjustment. In the *anchor filtering step*, Graphmap2 then uses a variation of the knapsack algorithm to find the optimal set of anchors and then uses KSW2 [11] aligner to perform piecewise affine gapped alignment between these anchors producing *first-phase* alignments. These alignments are then processed in the *Exon extending* and *Exon boundaries adjustment* steps which improve the quality of the *first-phase* alignments. Improved alignments are then split into exons which are further grouped and modified in *Exon grouping and adjustment* phase. Exons are grouped based on their position on the reference genome in order to group together exons transcribed from the same transcript. Exon groups are analysed and modified in *Exon offset calculation* and *Exon adjustment* steps. *Exon offset calculation* step calculates the correct start and end locations for all exons in the same group, for every group of exons. In *Exon adjustment* step incorrect exon’s starting or ending locations are adjusted to match previously calculated start and end locations of the whole group. *Exon adjustment* phase produces final alignments which are the output of the Graphmap2 tool.

**Figure 1.**
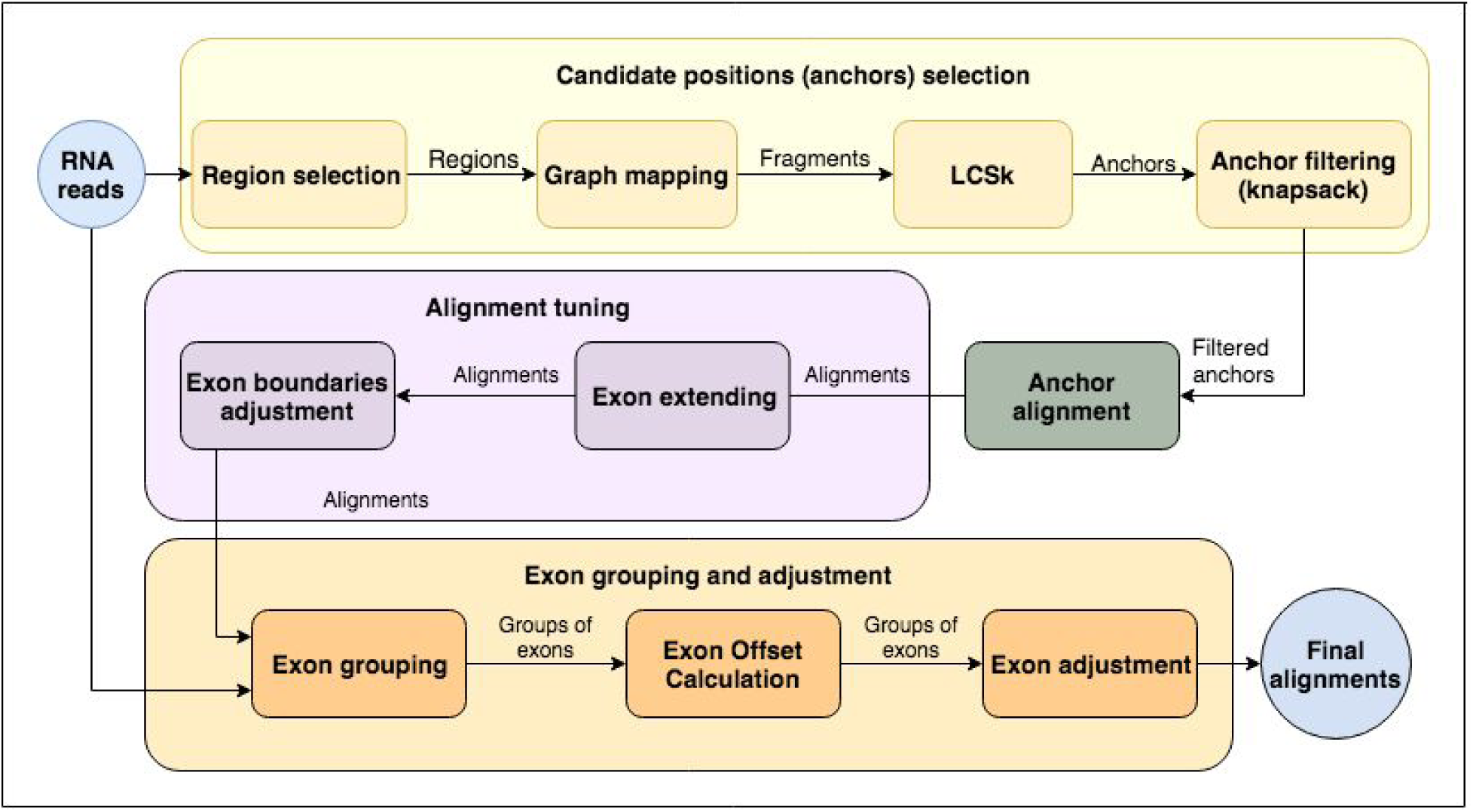
Graphmap2 read alignment process.

### Candidate positions selection

#### Anchor filtering

After completing the *LCSk* step of the Graphmap algorithm, produced anchors are then processed in order to obtain the optimal set of resulting anchors that are used to construct the alignment. If we denote reference starting and ending location in an anchor as *x*_*s*_ and *x*_*e*_ respectively, and query starting and ending location as *y*_*s*_ and *ye* respectively, we can consider every anchor as a two-dimensional line where the starting point of the line is *T* _*s*_(*x*_*s*_, *y*_*s*_) and ending point is *T* _*e*_(*x*_*e*_, *ye*). Also, every anchor has its fitness *f*, which in the simplest form is the number of bases *d* covered by the anchor. We can formalize the problem of finding the optimal set of anchors as follows: from the set *C* of *N* anchors, *C*_*i*_ = (*T* _*si*_, *T* _*ei*_, *d*_*i*_) ∈ *C*, we want to find an optimal set of *k* anchors

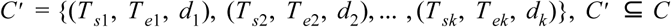

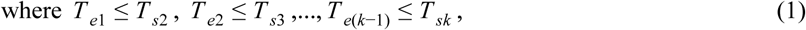

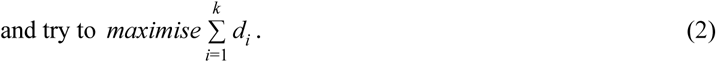

The problem defined in this way is equivalent to 0-1 knapsack problem and we can use the same algorithm to solve it. 0-1 knapsack problem can be described as: for a given set *E* of *N* elements, where every element *e* has its weight *w* and fitness *f*, and with a limit on maximum weight T, we need to find *E*′ ⊆ *E* whose sum of weights is not greater than T, and whose sum of fitnesses has maximum value, or formally:

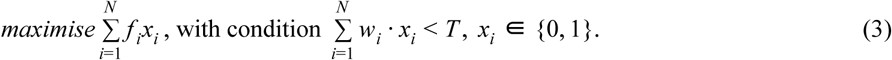

For the current set of anchors, the weight of the element is the length of query in the anchor |*ye* − *y*_*s*_|, while the fitness of the anchor is the number of covered bases *d*. The total weight T is the total length of the processed read. By solving this optimisation problem and finding *x*_*i*..*N*_ ∈ {0, 1} we find the resulting set of anchors that construct the reads’ alignment. Since Graphmap maps reads to both the original reference sequence and its reverse complement, this algorithm finds the optimal set of anchors regardless of the strand of the alignment.

With this algorithm we expect to find the optimal set of anchors which are used to generate the alignment of the read, regardless of their distance on the reference i.e. the gap between the anchors. The process of alignment is performed using the KSW2 aligner which aligns two adjacent anchors with a possible gap between them. For two adjacent anchors *C*_*i*_ = (*T* _*si*_(*x*_*si*_, *y*_*si*_), *T* _*ei*_(*x*_*ei*_, *y*_*ei*_)) and *C*_*i*+1_ = (*T _*s*(*i*+1)_(*x*_*s*(*i*+1)_, *y*_*s*(*i*+1)_*), *T* _*e*(*i*+1)_(*x*_*e*(*i*+1)_, *y*_*e*(*i*+1)_)) the alignment *a*_*i*_ is generated with KSW2 for the subset of read [*y*_*si*_, *y*_*e*(*I*+1)_] and the subset of reference [*x*_*si*_, *x*_*e*(*I*+1)_]. The portion of the alignment *a*_*i*_ that corresponds to the subset of read [*y*_*si*_, *y*_*s*(*i*+1)_] is then extracted and appended to the resulting alignment.

Because of high error rate generated by third generation sequencers, the optimal set of anchors can still fail to cover all of the exons of the read or can often have errors at the ends of gaps in the alignment. This is why we introduce several methods that try to improve alignments generated by the anchor alignment step. They are described in the following chapters.

### Alignment tuning

#### Exon extending

Because a different set of exons can identify different gene expression, it is important that alignment of RNA reads contains all exons from that read correctly identified. Since some alignments tend to lose an exon because of errors at edges of the read, we have implemented a tuning method that tries to align clipped ending of a read to the reference. The clipped part of the read is aligned to reference and if the alignment of that clipped part of the read contains no more than 15% of deletions, insertions or substitutions, it is considered as a new found exon and it is appended to the alignment. Before aligning clipped parts of the reads, poly(A) tail identification is conducted first. If the clipped part of the read is identified as a poly(A) sequence, it is not further analyzed.

Algorithm 1 shows the process of improving an alignment by aligning edges of reads that have been marked as clipped after *Anchor alignment* step. The unaligned portions at the beginning or the end of a read are denoted as *unaligned edges*. If an *unaligned edge* is at the end of the read, we first check if the unaligned edge contains a poly A tail, and if so, the edge is not processed any further. After that, the reference region, with length *windowLength*, whose one end corresponds to the end of the edge of the read is found. The *windowLength* was experimentally set to 8000. The edge of the read is then aligned to the reference region and the resulting alignment, denoted as *edge alignment* is processed further. The first occurred exon (its alignment) found in the edge alignment is extracted, and if its length is greater than the kmer length (original Graphmap parameter) and its alignment identity is at least 85%, it is appended to the original alignment. If the found exon is not on the starting position of the edge alignment, a gap, of length equal to the starting position of the exon in the edge alignment, is added to the original alignment.

##### Algorithm 1 Extend alignment

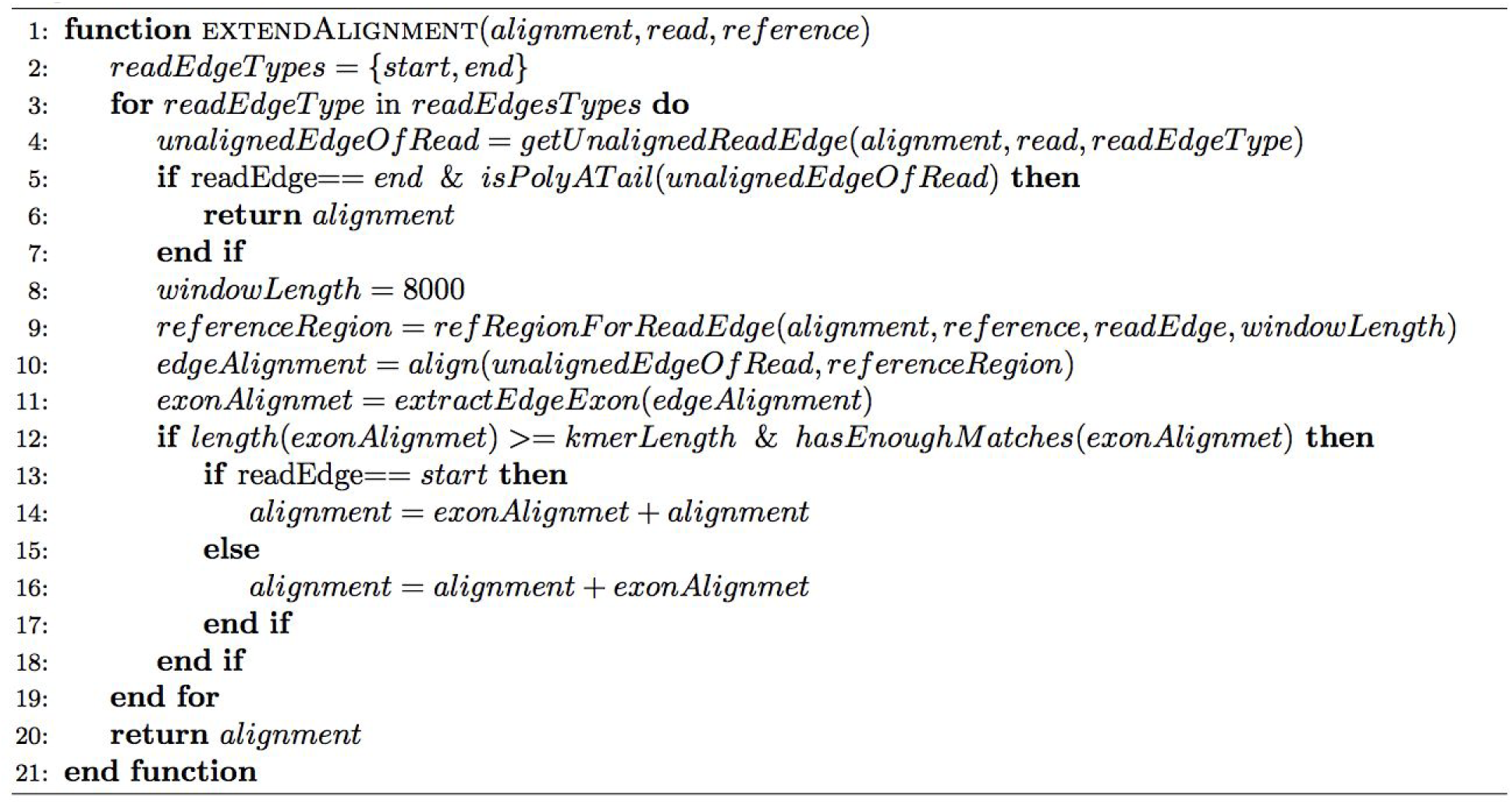

#### Exon boundaries adjustment

Because single base precision is important for RNA mapping, we have implemented an algorithm the improves exon boundaries using the information about donor-acceptor splice sites. Introns usually have two distinct nucleotides at either end: GT at the 5’ end (CT for reversed strand) and AG at the 3’ end (AC for reversed strand). These nucleotides are a part of splicing sites. Since there is a high probability that introns should begin and end with these bases, for every read with spliced alignment, every two neighbouring exons *e*_*i*_ *and e*_(*i*+1)_ in the alignment are analysed. If bases GT (CT) are found somewhere around the end of exon *e*_*i*_, up to 5 bases left or right on the reference, and if bases AG (AC) are found somewhere around the start of the exon *e*_(*i*+1)_, up to 5 bases left or right on the reference, the alignment is modified so that exon *e*_*i*_ ends exactly next to AG (AC) and exon *e*_(*i*+1)_ starts exactly after GT (CT) splice sites. This procedure is presented in Algorithm 2.

##### Algorithm 2 Exon boundaries adjustments

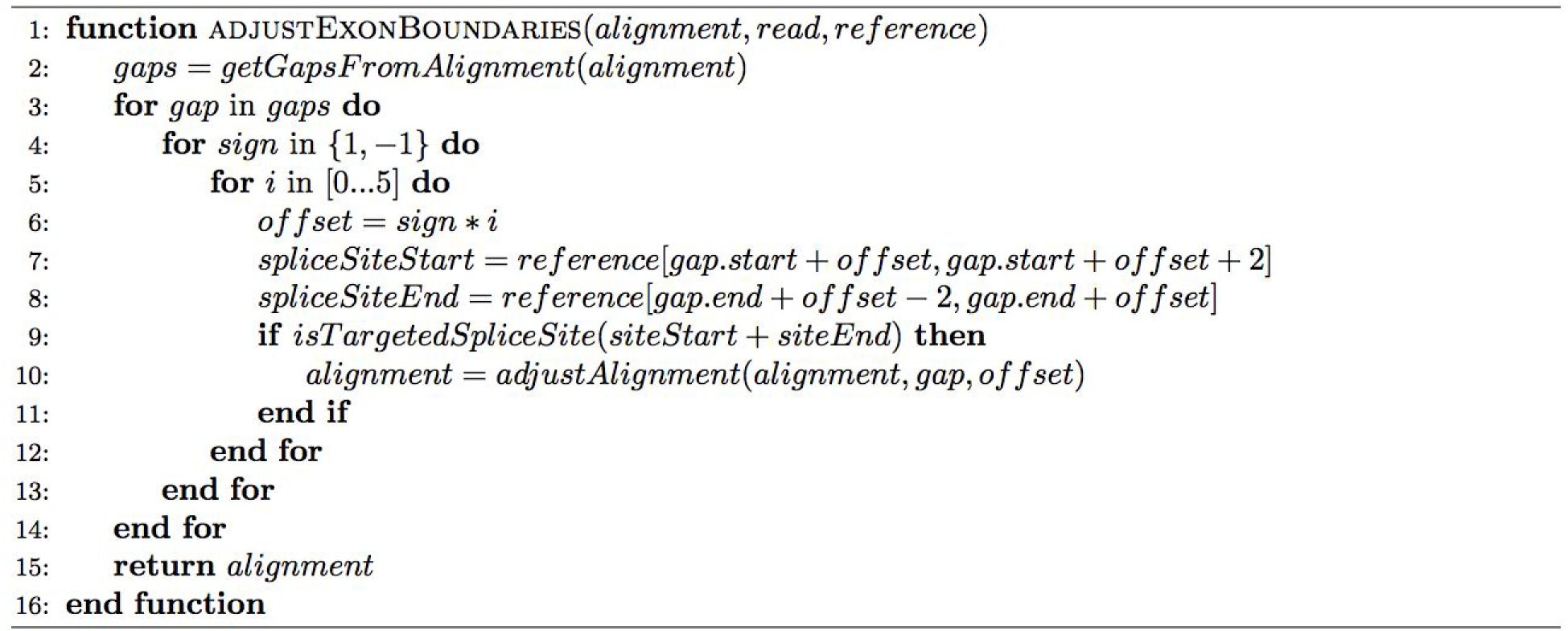

### Exon grouping and adjustment

#### Exon grouping

As already mentioned, *first-phase* alignments are created from anchors using KSW2. The most common errors that occur in alignments produced by the *anchor alignment step* are: (1) false exon, (2) missed exon and (3) incorrect intron ends. False exon error occurs when two parts of a read that belong to a single exon and should be aligned together are aligned separately. Missed exon error is an error where two parts of a read that belong to separate exons and should be aligned separately, are aligned together. Wrong intron ends error occurs when a part of a read belonging to one exon is aligned as the edge of another exon. These errors usually occur due to sequencing errors.

##### Algorithm 3 Group exons

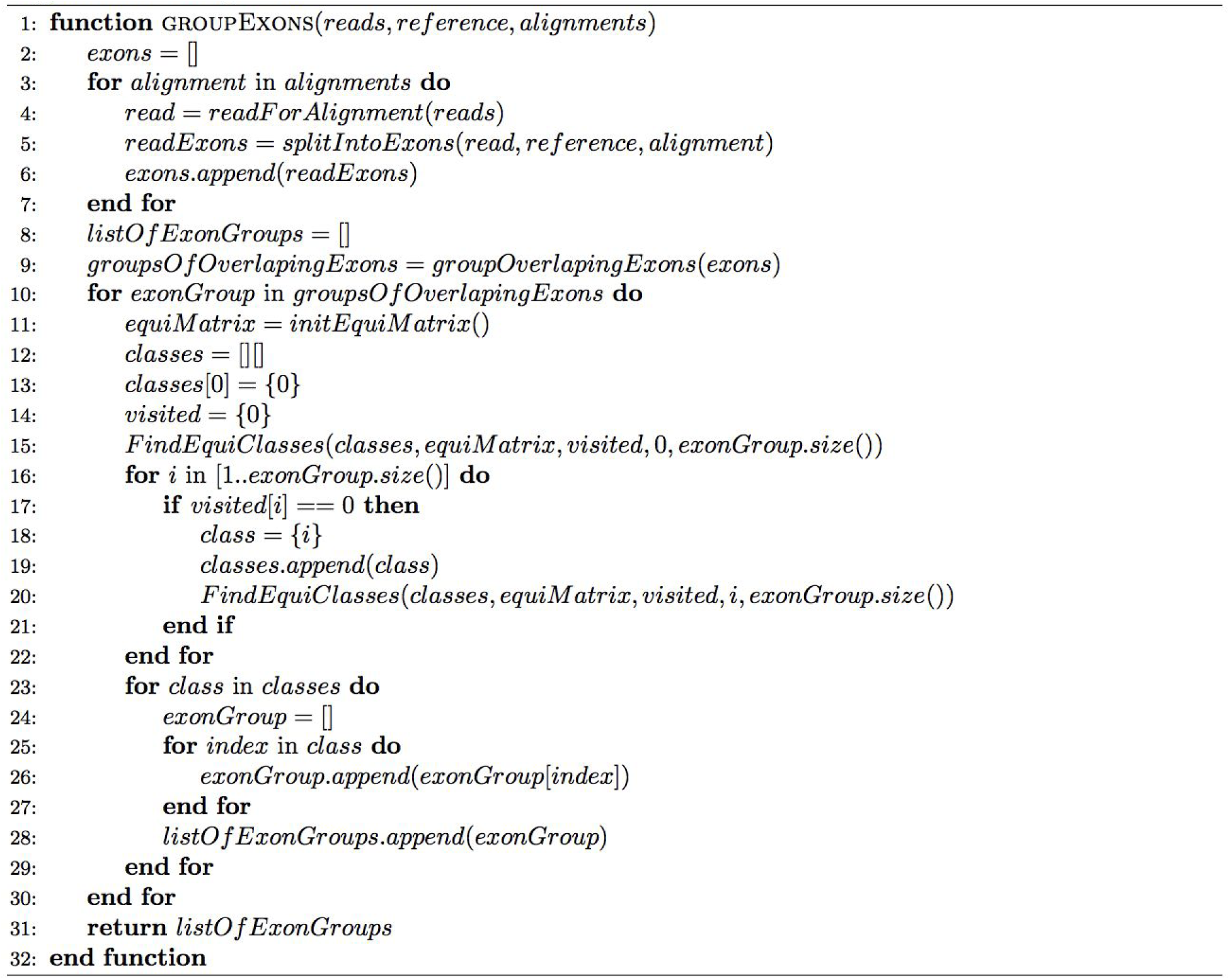

The *exon grouping and adjustment* phase of the Graphmap2 tries to overcome these errors by improving on already calculated alignments. Since it is expected that a number of reads is transcribed from the same region on the reference, some of them might have previously described errors, while others might be correctly aligned. *Exon grouping* step starts by identifying groups of reads that cover the same transcript location in the reference.

Reads are placed in the same group if they mutually overlap. Ideally, all the reads in the group should belong to the same transcription region, but this is not always the case. This is why every group is analysed using coverage information for the reads in the group. The coverage information of the group is calculated from their alignments. For every group of alignments we denote the coverage of a single base in the reference as the number of alignments that contain that reference base. The average coverage of the group is the total number of coverage values of all bases in the reference that are contained in alignments of the group, divided by the number of those bases. If all the alignments in the group are identical, then the average coverage of the group would be equal to the number of reads in the group. The more alignments contain different reference bases, the lower the average coverage of the group. The groups with the average coverage of the group greater than 50% of the number of reads in the group are chosen for the *Exon adjustment* phase.

For every group of reads their alignments are split into exons by splitting cigar strings by gapped components. Every exon is modeled with: (1) start and stop location on the reference, (2) read id, (3) index of exon in the alignment, (4) cigar string of the exon, (5) *start offset* by which the start location of the exon should be modified and (6) *end offset* by which the end location of the exon should be modified. These exons are then grouped as shown in algorithm 3.

We define the relation:

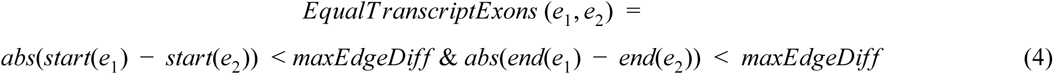

*e*_1_, *e*_2_ ∈ *E*_*group*_ where group. *E*_*group*_ is set of exons for one group of reads and *e*_1_ and *e*_2_ are two exons from the same group

##### Algorithm 4 Initilize Equivalency Matrix

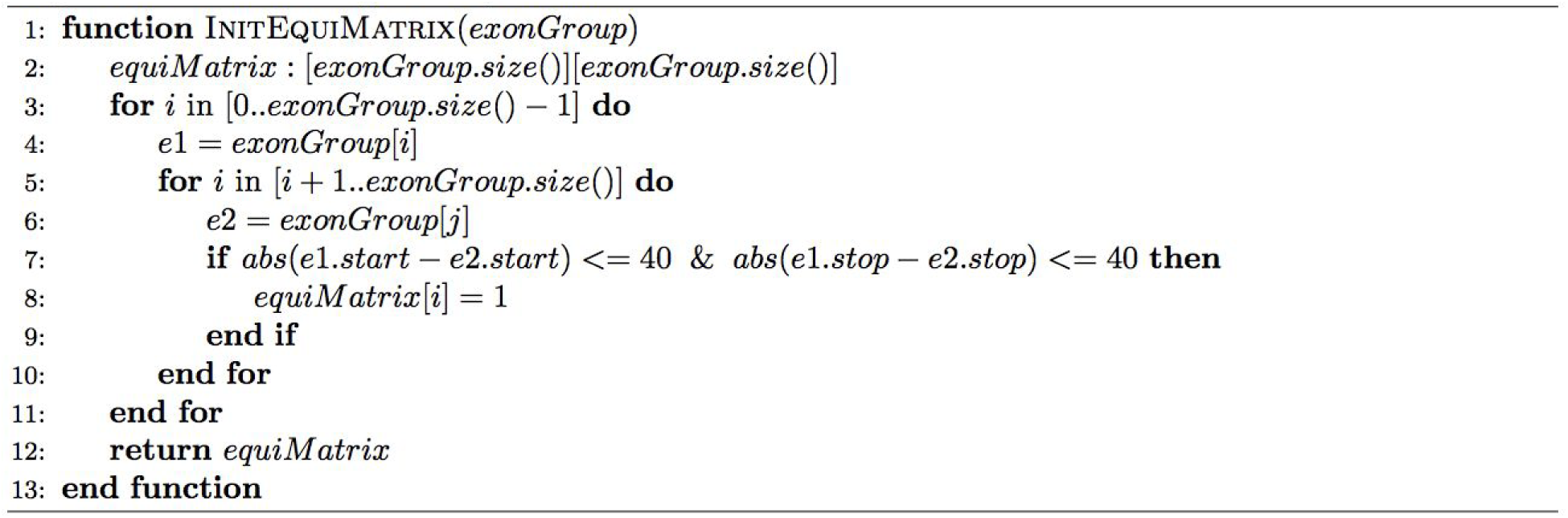

Two exons whose start and end location don’t differ more than *maxEdgeDiff* number of bases are in *EqualTranscriptExons* relation and we consider them to be transcribed from the same transcript region. The algorithm 3 finds all equivalence classes of relation *EqualT ranscriptExons* thus finding all equivalence groups of *E*_*group*_ exons. It uses equivalence matrix which is a *EqualT ranscriptExons* relation matrix and is initialized as shown in algorithm 4. Parameter *maxEdgeDiff* was experimentally set to 40.

Using equivalency matrix, we recursively find all groups of exons for current group of reads using algorithm 5. Every group of exons is then analysed in the *Exon edge offsets calculation* step.

##### Algorithm 5 Find Equivalent Classes

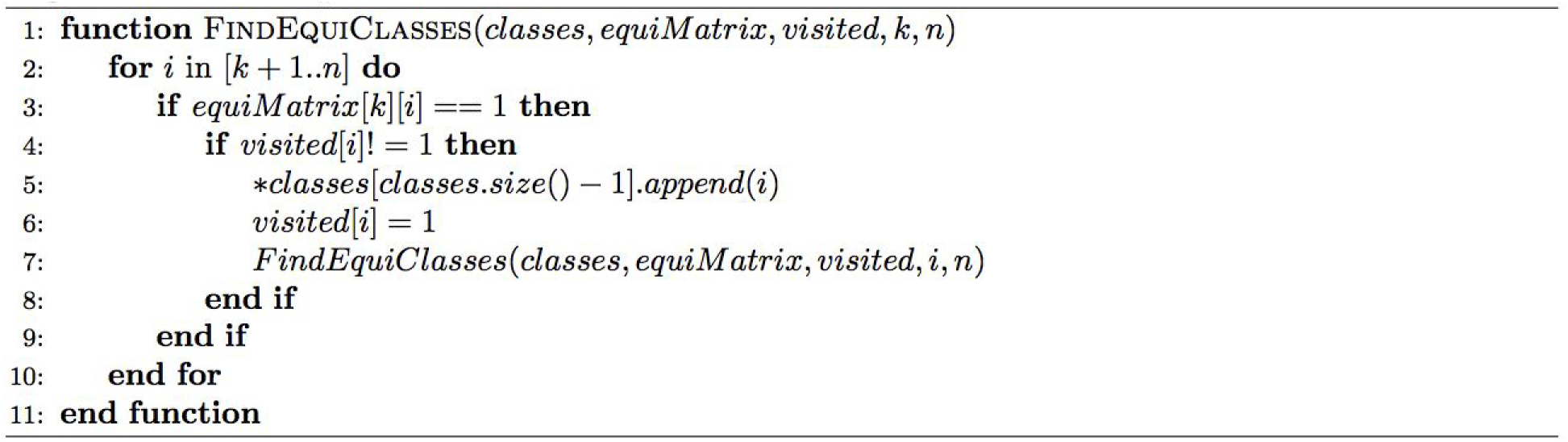

#### Exon edge offsets calculation

Because we assume that the exons in the same group are transcribed from the same region in the reference, we can also assume that all of the exons in the group should have identical start and end locations. Correctly aligned exons in a group facilitate correct identification of start and end location for the whole group. Exons from the group that have alignment errors and do not have start or end location identical to group’s start or end location are then corrected by modifying their start or end locations in the *Exon adjustment* step.

##### Algorithm 6 Find Exons Offsets

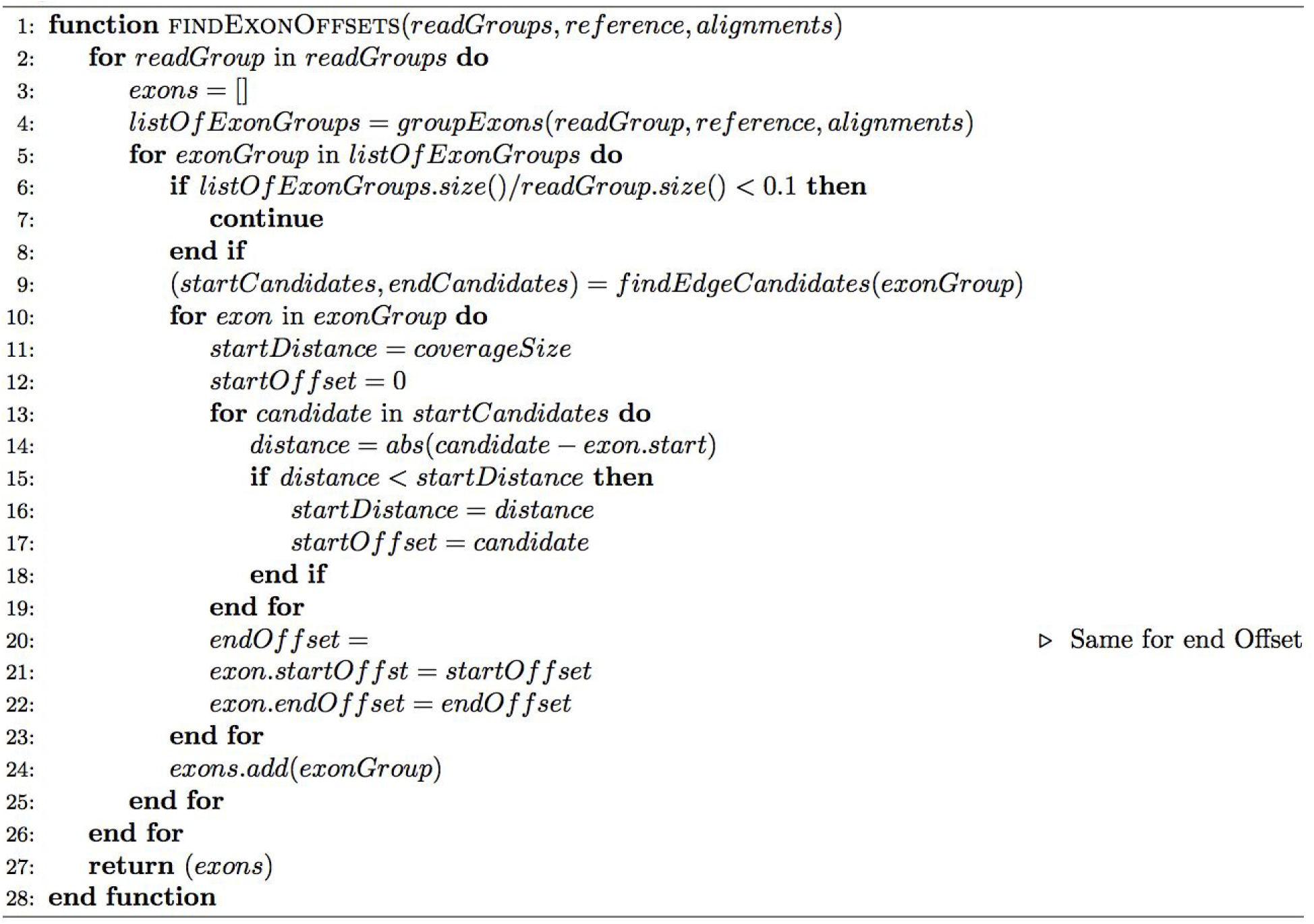

The process of determining group’s start and end location is shown in algorithms 6 and 7. For every group of exons that contains exons from at least 10% of reads in the analysed group of reads, the candidate start and end locations for whole group are determined. As shown in algorithm 7, for some exon group two coverage arrays are constructed using the following procedure:

##### Algorithm 7 Find Exon Edges Candidates

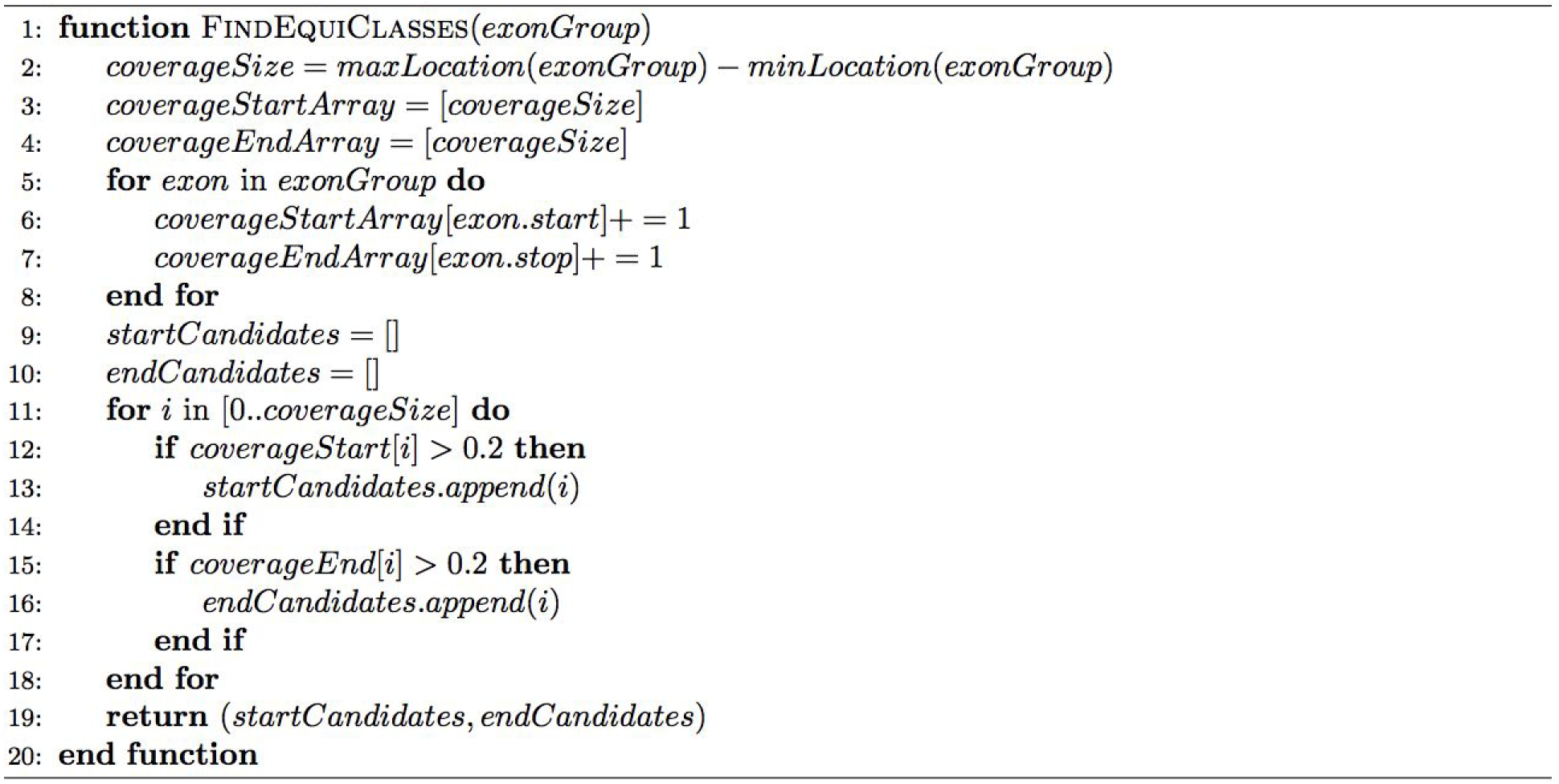

for every exon *e*_*i*_ ∈ *E* in the group of exons *g*_*k*_ ∈ *G* of group of reads R where *i* ∈ {1… *m*} and |*E*| = *m*, and |*R*| = *l* we denote

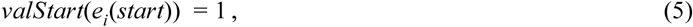

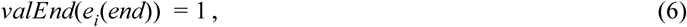

for every location *j* ∈ {*s*, … *e*} in the reference genome, where *s* is the start location on the reference of the left most exon in the group, and *e* is the end location on the reference of the most right exon in the group.

The start and end coverage of the location *j* in the group of exons is defined as:

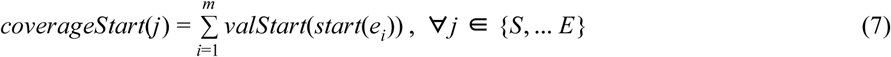

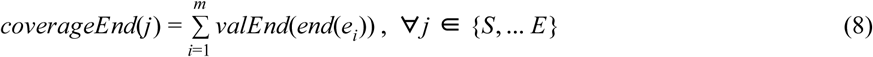

From these values start and end coverage arrays are constructed and used to determine candidates for start and end locations of all exons in the group. All locations in these coverage arrays that satisfy

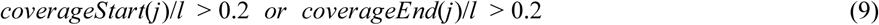

are chosen as exon start and end location candidates. Multiple candidates are chosen because certain groups can contain exons that are transcribed from more than one transcript with similar, but not identical start and end locations on the reference. This is why every group is assigned a set of possible candidates of start and end locations for the whole group.

Finally, as shown in algorithm 6, using start and end location candidates, start and end offsets are calculated for every exon in the group. These offsets represent the number of bases that need to be added or removed from the start/end of the exon so that its start/end is equal to one of the start/end candidate locations previously found for the whole exon group. A start/end candidate location, with the shortest distance to the current exon’s start/end location, is chosen as the desired start/end location for that exon and is used to calculate start/end offset for that exon which is then used in *Exon adjustment* step to modify exons start and end location.

#### Exon adjustment

Figure 2 shows the simplified process of adjusting the alignments of two neighbouring exons using *left offset* value and *right offset* value calculated in the previous *exon edge offsets calculation* step. More specifically, right edge of the left observed exon and left edge of the right observed exon are adjusted. Reference components of the two observed exons are merged omitting the gap between the exons. Left exon’s reference component is trimmed or extended on it’s right edge by *right offset* number of bases, and right exon’s reference component is trimmed or extended on it’s left edge by *left offset* number of bases. The resulting merged reference component is then aligned with the read component, created by simply merging both read components of the two observed exons. The resulting alignment is then split into two parts using *left exon desired reference length* which is the previous length of left exon’s reference component subtracted by that exon’s *right offset* value. The left part of the alignment is cut so that it’s reference component has *left exon desired reference length* and it is used as the left exon alignment.

##### Algorithm 8 Adjust Read Exons

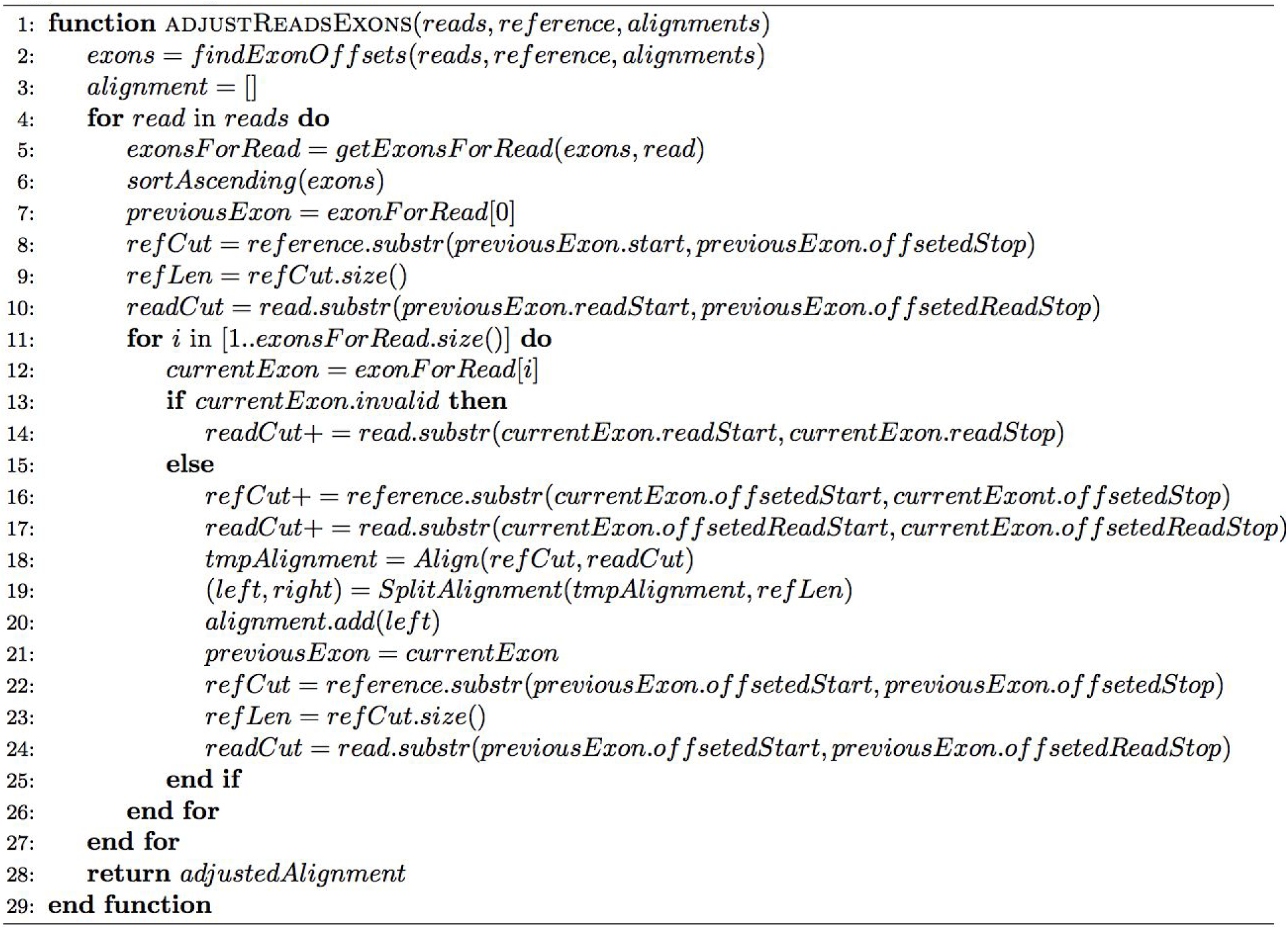

**Figure 2.**
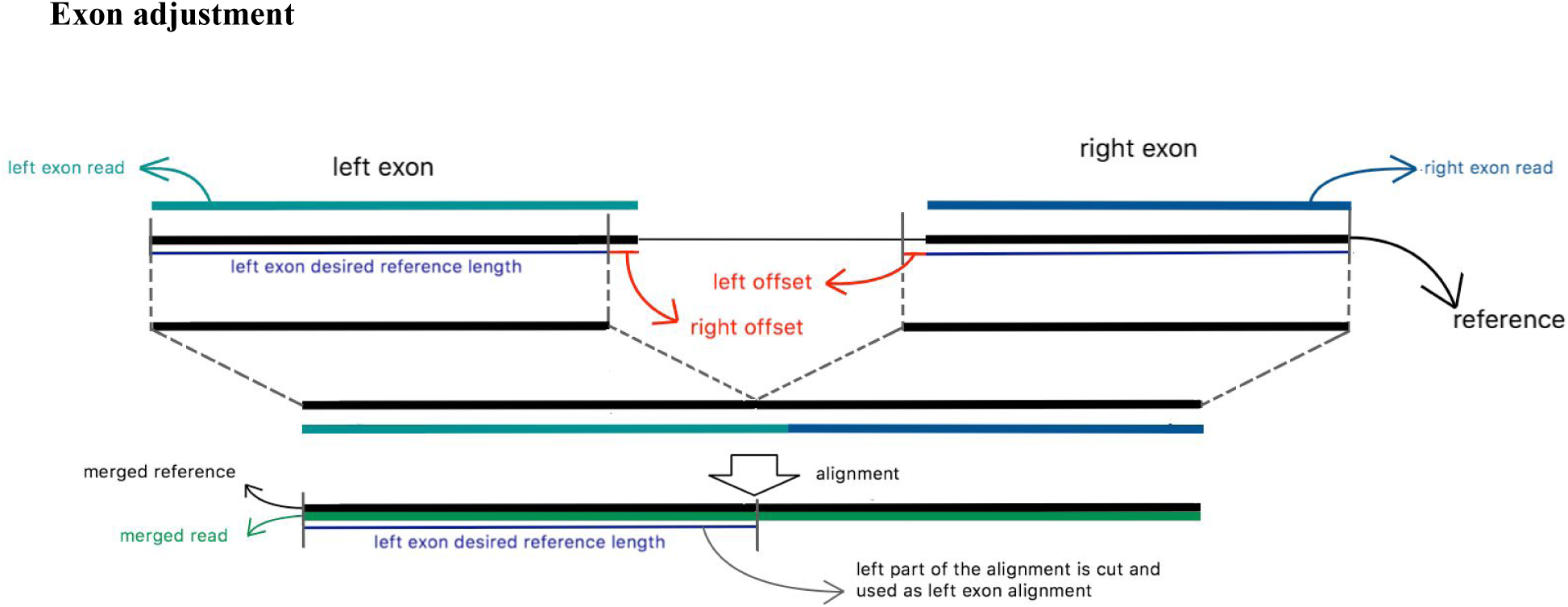
Exon edges adjustment.

Algorithm 8 shows the general outline of the exon adjustment implementation. With complete *exon grouping and adjustment* procedure, a great number of reads is improved by using the fact that high number of previously generated alignments is correctly aligned and can be used to improve same-transcript reads with alignment errors.

#### Alignment scoring

When making alignment modifications in *Alignment tuning* phase or *Exon grouping and modification* phase, the resulting alignments are compared to the alignments from the previous phase. When comparing two alignments, for both alignments, the E-value is calculated with parameters used in [7]: match = 5, mismatch = −4, gapopen = −8 and gapextend = −6, and, if the difference between two E-values is greater than 0.05, or if the read is not spliced, the alignment with the greater E-value is chosen as the final alignment. If the absolute difference between E-values of two spliced alignments is less than 0.05, which means that two alignments are of almost equal quality, the quality of the edges of the exons in the alignments is further analysed. Because in most cases alignments’ errors are located at the edges of exons, the alignments’ exon edges are scored by calculating matches values with parameters: match = 5, any other error = −4. The edge of an exon has experimentally been set to length 10. The higher scored alignment is chosen as the final alignment of the read.

All the improvements described in this chapter affect alignments of reads that are usually close to being fully correct or miss one or two exons. By applying these improvements we achieve high precision and sensitivity of the Graphmap2 tool, which outperforms other existing RNA mapping tools on majority of datasets. The evaluation of the Graphmap2 performance is given in the next chapter.

## III. RESULTS

Graphmap2 was compared to two best performing RNA seq mappers evaluated in [3], Minimap2 and GMAP. Minimap2 was downloaded from https://github.com/lh3/minimap2 (v2.17), while GMAP (version 2019-06-10) was downloaded from http://research-pub.gene.com/gmap/. All three tools were evaluated on 7 different datasets which contain reads sequenced by third generation sequencers. Four datasets are synthetic, created from the following organisms: (1) Saccharomyces cerevisiae S288 (baker’s yeast), (2) Drosophila melanogaster r6 (fruit fly) and (3) Homo Sapiens GRCh38.p7 (human). Three synthetic datasets simulate reads generated by PacBio sequencers, while the fourth dataset simulates reads generated by ONT sequencers. Synthetic datasets are the ones used in [4]. Remaining three datasets are real datasets of Drosophila melanogaster, with two of them being produced by PacBio sequencers and one produced by ONT sequencers.

When finding the origin of transcription of RNA reads by aligning them to a reference genome, there are two main goals that need to be achieved: (1) generated alignment needs to be precise up to a single nucleotide base, (2) all of the exons of the RNA read need to be found. The first goal is important because a miss of a single nucleotide can implicate a completely different protein coded from that RNA molecule, while not meeting the second goal can lead to misinterpretation of the gene expressed by mapped RNA molecule. To measure the quality of achieving these goals the alignments generated by all three tools were compared to annotations used in [3] where the starting and ending positions of each exon in the alignment were compared to starting and ending positions of exons in the annotation. The reads whose alignment has the same number of exons as the corresponding annotation and has all of its exons overlapped with the corresponding exons in the annotation by at least one nucleotide, were denoted as *hit-all* reads. Furthermore, the reads that are denoted as *hit-all* read, but also satisfied next condition:

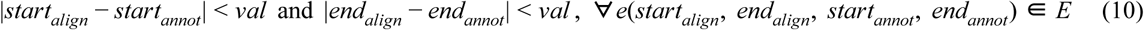

were denoted as *correct* reads, where the *E* is the set of exons in the alignment, *start*_*align*_ and *end*_*align*_ are the starting and ending reference positions of the exon *e* in the alignment and *start*_*annot*_ and *end*_*annot*_ are the starting and ending reference positions of the exon *e* in the corresponding annotation. The specific *correct* measure depends on the chosen value *val*. In this paper we present the evaluation of the tools using two *correct* measures: (1) *correct-0* for *val* = 0, and (2) *correct-5* for *val* = 5. The first, *correct-0* measure is used to evaluate how well tools find reads that are fully correct, up to a single nucleotide, while the second measure, *correct-5*, is used to evaluate if the tools align reads correctly but are not completely precise, with a few bases off. For three measures defined previously: *hit-all, correct-0* and *correct-5*, the precision is calculated as the number of reads satisfying the measure divided by the number of aligned reads and recall is calculated as the number of reads satisfying the measure divided by the total number of reads. Finally, the F-value is calculated as

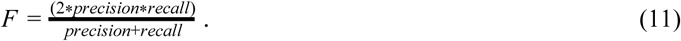

Table 1 shows the result of the evaluation on all seven datasets with their F-value for *correct-0* measure and *hit-all* measure results. Hit-all measure was calculated only for synthetic datasets because for real datasets we cannot know the exact origin of the read and cannot determine the exons from the associated annotation that need to be hit by the read’s alignment. We can see that Graphmap2 is the most sensitive tool and has the best results in both *correct-0* measure across four simulated datasets. Because of the strict condition for an alignment to be *correct-0* (0-bp distance from the expected location) the values of *correct-0* measure are low across all datasets, but Graphmap2 is the only tool that has recall for *correct-0* measure above 30% for 6 out of 7 datasets. Supplementary Table 1 shows that Graphmap2 also achieves the best results for correct-5 measure, having the highest number of reads aligned with almost fully correct alignments. Except for the GMAP results for dataset2, all the tools have *hit-all* measure higher than 70%. This is why there are no great differences between *hit-all* measure results. Still, Graphmap has the best precision and recall results for *hit-all* measure across all datasets, finding the highest number of alignments with all exons hit. We can see that the Graphmap2 is the only tool that consistently achieves good results in *correct-0* and *hit-all* measures at the same time, thus the most consistently maps reads correctly and with all exons hit at the same time.

**Table 1.**
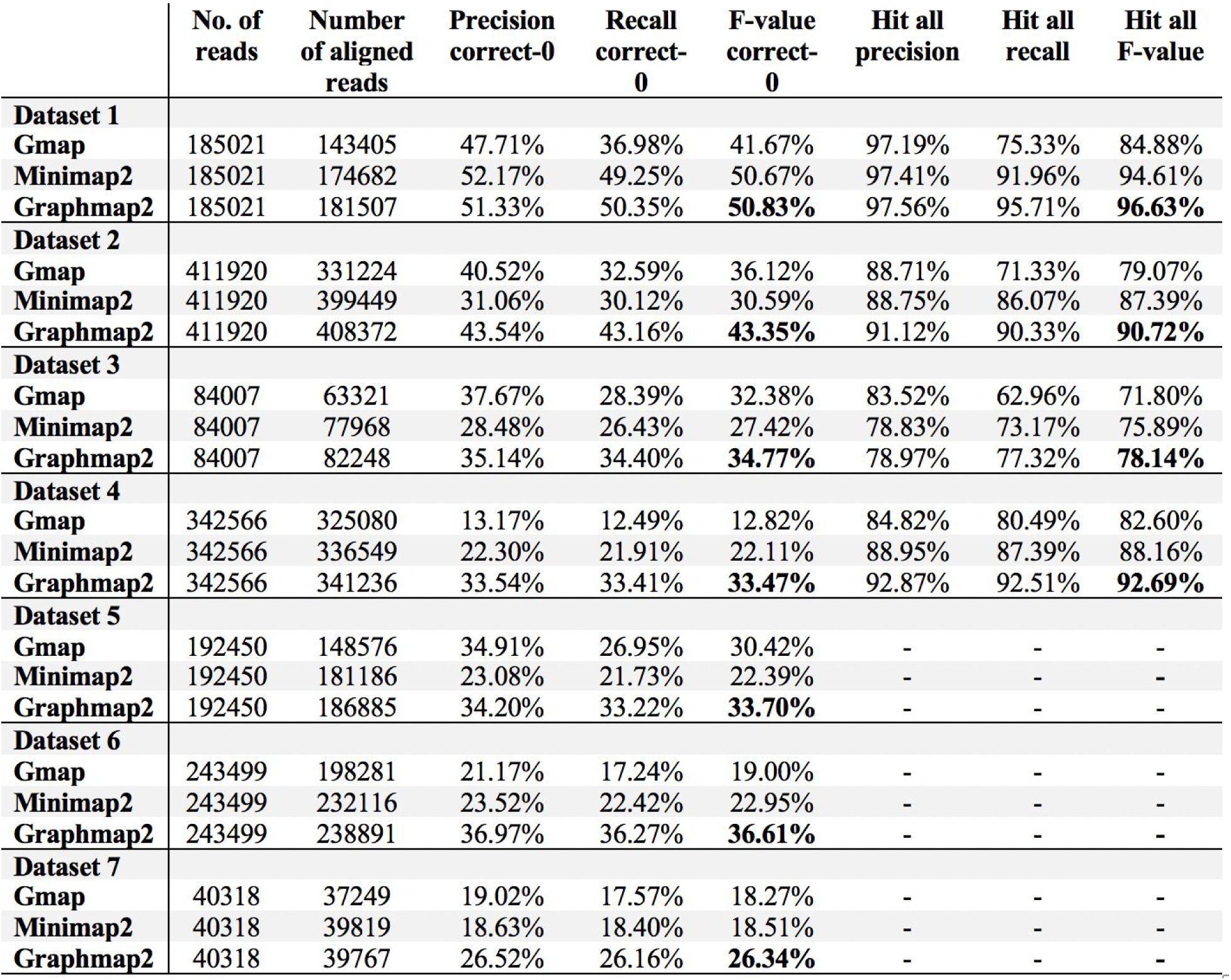
Results of evaluation of three best performing RNA-seq mapping tools.

|  | No. of reads | Number of aligned reads | Precision correct-0 | Recall correct-0 | F-value correct-0 | Hit all precision | Hit all recall | Hit all F-value |
| --- | --- | --- | --- | --- | --- | --- | --- | --- |
| **Dataset 1** |  |  |  |  |  |  |  |  |
| **Gmap** | 185021 | 143405 | 47.71% | 36.98% | 41.67% | 97.19% | 75.33% | 84.88% |
| **Minimap2** | 185021 | 174682 | 52.17% | 49.25% | 50.67% | 97.41% | 91.96% | 94.61% |
| **Graphmap2** | 185021 | 181507 | 51.33% | 50.35% | **50.83%** | 97.56% | 95.71% | **96.63%** |
| **Dataset 2** |  |  |  |  |  |  |  |  |
| **Gmap** | 411920 | 331224 | 40.52% | 32.59% | 36.12% | 88.71% | 71.33% | 79.07% |
| **Minimap2** | 411920 | 399449 | 31.06% | 30.12% | 30.59% | 88.75% | 86.07% | 87.39% |
| **Graphmap2** | 411920 | 408372 | 43.54% | 43.16% | **43.35%** | 91.12% | 90.33% | **90.72%** |
| **Dataset 3** |  |  |  |  |  |  |  |  |
| **Gmap** | 84007 | 63321 | 37.67% | 28.39% | 32.38% | 83.52% | 62.96% | 71.80% |
| **Minimap2** | 84007 | 77968 | 28.48% | 26.43% | 27.42% | 78.83% | 73.17% | 75.89% |
| **Graphmap2** | 84007 | 82248 | 35.14% | 34.40% | **34.77%** | 78.97% | 77.32% | **78.14%** |
| **Dataset 4** |  |  |  |  |  |  |  |  |
| **Gmap** | 342566 | 325080 | 13.17% | 12.49% | 12.82% | 84.82% | 80.49% | 82.60% |
| **Minimap2** | 342566 | 336549 | 22.30% | 21.91% | 22.11% | 88.95% | 87.39% | 88.16% |
| **Graphmap2** | 342566 | 341236 | 33.54% | 33.41% | **33.47%** | 92.87% | 92.51% | **92.69%** |
| **Dataset 5** |  |  |  |  |  |  |  |  |
| **Gmap** | 192450 | 148576 | 34.91% | 26.95% | 30.42% | - | - | - |
| **Minimap2** | 192450 | 181186 | 23.08% | 21.73% | 22.39% | - | - | - |
| **Graphmap2** | 192450 | 186885 | 34.20% | 33.22% | **33.70%** | - | - | - |
| **Dataset 6** |  |  |  |  |  |  |  |  |
| **Gmap** | 243499 | 198281 | 21.17% | 17.24% | 19.00% | - | - | - |
| **Minimap2** | 243499 | 232116 | 23.52% | 22.42% | 22.95% | - | - | - |
| **Graphmap2** | 243499 | 238891 | 36.97% | 36.27% | **36.61%** | - | - | - |
| **Dataset 7** |  |  |  |  |  |  |  |  |
| **Gmap** | 40318 | 37249 | 19.02% | 17.57% | 18.27% | - | - | - |
| **Minimap2** | 40318 | 39819 | 18.63% | 18.40% | 18.51% | - | - | - |
| **Graphmap2** | 40318 | 39767 | 26.52% | 26.16% | **26.34%** | - | - | - |

Table 2 shows the execution time in seconds for the three tools for every tested dataset. The testing was done by executing tools using 12 threads on a computer with 12 processor cores. It is clear that Minimap2 is the fastest tool (9 to 40x faster than Graphmap2). GMAP shows to be the slowest tool being 3-7 times slower than Graphmap2. Since Graphmap2 achieves high sensitivity by mapping reads in three phases and executes *Exon extending* and *Exon adjustment* steps which also calculates exact alignment of read parts, it is expected not to be as fast as Minimap2. Graphmap2s’ speed greatly suffers because of its’ high sensitivity which was the main focus of Graphmap2 development. However, there is always room for further research in how to improve Graphmap2s’ speed while retaining the sensitivity. As correctness of sequenced reads improves, a stricter anchor filtering could potentially lead to faster execution time.

**Table 2.** Execution time in seconds.

| Tools | Dataset 1 | Dataset 2 | Dataset 3 | Dataset 4 | Dataset 5 | Dataset 6 | Dataset 7 |
| --- | --- | --- | --- | --- | --- | --- | --- |
| **GMAP** | 6675 | 11093 | 1334 | 13535 | 11615 | 15945 | 2012 |
| **Minimap2** | 24 | 142 | 45 | 140 | 186 | 146 | 28 |
| **Graphmap2** | 904 | 3242 | 488 | 3411 | 1946 | 1783 | 262 |

## IV. PREVIOUSLY UNKNOWN GENE IDENTIFICATION

In this chapter we present the analysis of Graphmap2’s ability to find mappings of yet unidentified genes. To confirm the existence of new found genes, we analysed alignments produced by the three tools used in the evaluation. In this analysis we used the three real datasets and the alignments that didn’t match any known annotation. To remove as much coincidence as possible, this analysis was done highly conservatively and the alignments that did not overlap along at least 50% of their length with any other alignment were also left out of the analysis. This was done because the mapping of at least two reads onto the same region on the reference suggests that that region could be a new gene and not a coincidence. The alignments used in this analysis are denoted as gene-candidate alignments. Supplementary Table 2 shows the statistics of these alignments for three tools and three real datasets: (1) total number of reads with gene-candidate alignments, (2) number of spliced alignments, (3) number of spliced alignments containing at least one pair of two neighbouring exons *e*_*i*_ *and e*_(*i*+1)_, denoted as *gene-identifying exons* where *e*_*i*_ exon ends right next to GT (CT) splice site and *e*_(*i*+1)_ exon starts right after AG (AC) splice site, (4) total number of neighbouring exons in the alignments and (5) total number of introns contained among gene-candidate alignments. Around 1000 reads for datasets 5 and 6 and more than 5000 reads for dataset 7 were found to have gene-candidate alignments, with 25%-50% of those being spliced alignments. These spliced alignments are important because they can be used in analysis of donor and acceptor splice sites at intron ends. Between 40% and 45% of spliced gene-candidate alignments have at least one pair of gene-identifying exons which is important evidence supporting the claim that gene-candidate alignments are correctly mapped to reference regions representing new found genes.

Gene-candidate alignments were further analyzed. Since we presume that overlapping reads identify the same gene, we construct a gene-candidate region as a union of mutually overlapping alignments, grouped similarly as in Graphmap2’s *Exon grouping* step. We analysed gene-candidate regions that contained at least two reads and had at least one read whose alignment contained at least one pair of gene-identifying exons.

Figure 3 shows the overlapping of gene-candidate regions with at least one pair of gene-identifying exons across three real datasets for all three tools. Graphmap2 has the highest number of gene-identifying regions found in all three datasets. These values show that even though datasets were sequenced with different sequencing technologies they contain reads whose alignments map on the same regions which are yet not found in annotations, furthermore strengthening the idea that newfound genes can be identified by Graphmap2.

**Figure 3.**
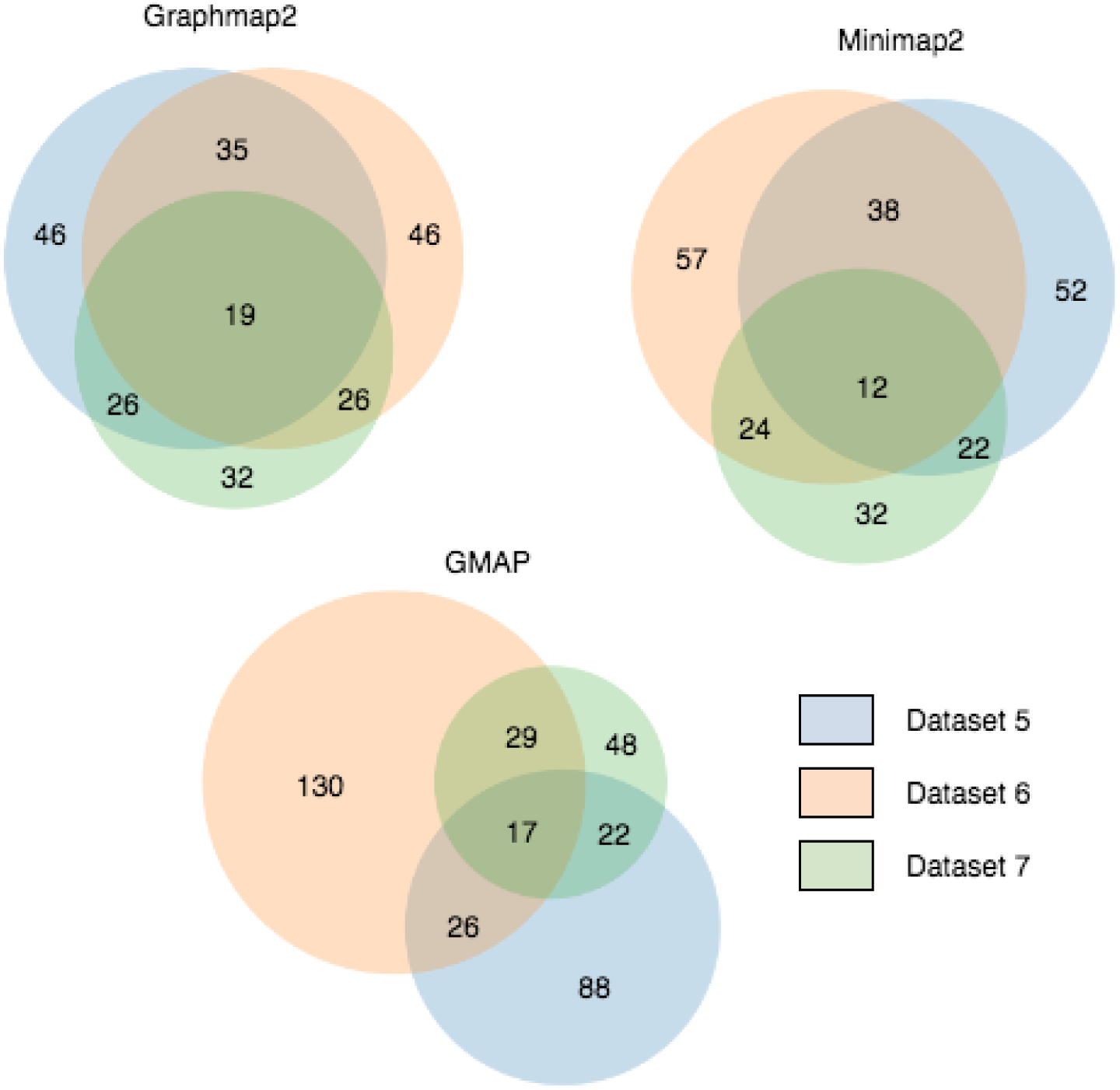
Overlapping of reads whose alignments did not hit an annotation among tools.

Figure 4 shows the number of these regions for three real datasets and the overlapping of regions found by three tools used in analysis. We can see that tools identified between 59 to 71 gene-candidate regions for dataset 5, 74 to 116 regions for dataset 6 and 32 to 36 regions for dataset 7, with about 40% to 60% of regions found by at least two tools. GMAP identified the most regions that don’t overlap with any of the regions produced by Minimap2 and Graphmap2 which suggest that GMAP produces the highest number of false positive regions, especially on datasets 6 and 7. Graphmap2 and Minimap2 have higher number of overlapping gene-candidate regions than either tool has with GMAP for datasets 5 and 6 which suggest that these two tools might identify previously unknown genes with higher accuracy than GMAP. High number of overlapping gene-candidate regions with gene-identifying exons confirms the hypothesis that previously unknown gene can be detected using long reads and splice-aware mappers.

**Figure 4.**
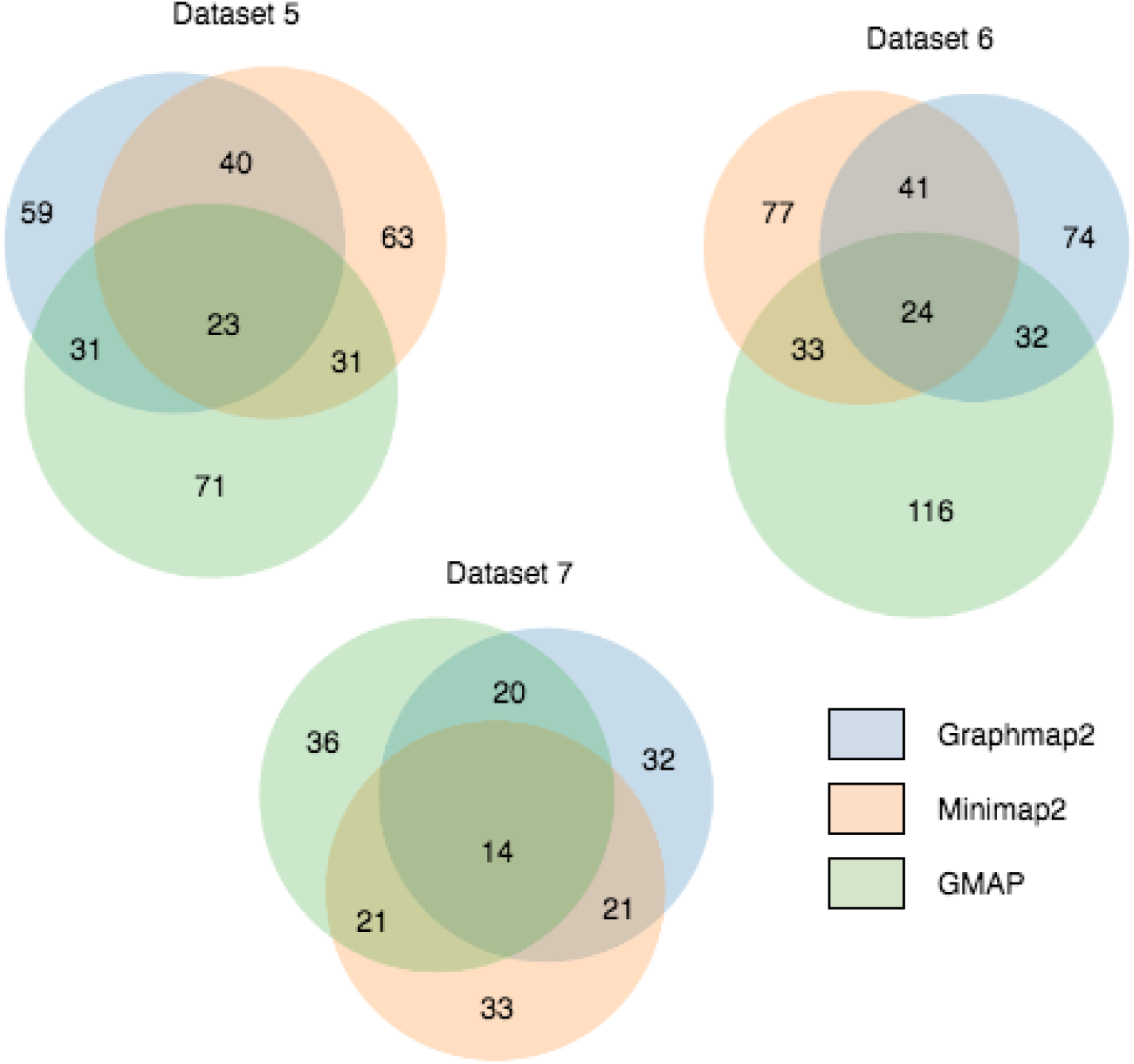
Overlapping of reads whose alignments did not hit an annotation among datasets.

## V. PREVIOUSLY UNKNOWN ISOFORM IDENTIFICATION

Aside from detecting new genes with no known annotations, we have also tried to discover new isoforms in regions where annotations are already known. Isoform detection was done using newest version of our evaluation tool RNAseqEval (https://github.com/lbcb-sci/RNAseqEval). During the regular evaluation process, a set of candidate annotations is constructed for each alignment, consisting of all annotations that overlap the alignment. From the set of candidate annotations, a *best_match_annotation* is chosen based on the number of nucleotides from the alignment that fall inside and outside of each candidate annotation. New isoforms are calculated only for those alignments that do not perfectly match the *best_match_annotation*. New isoforms are created using two methods: by combining existing annotations and by skipping small introns. Both methods are explained in detail on the RNAseqEval GitHub page. After all new isoforms are determined, they are evaluated, collected and mutually compared. Only isoforms that result in completely correct alignment and are supported by a minimum number of reads (currently set to 3) are reported.

The number of new isoforms discovered from alignments for each tool and dataset can be seen in Figure 5. We can see that Dataset 5, containing PacBio ROI reads with the lowest error rate, produces the highest number of new isoforms. Dataset 6 contains PacBio subreads with slightly higher error rate, resulting in a slightly lower number of new isoforms. Finally, Dataset 7 contains ONT reads with the highest error rate producing by far the smallest number of new isoforms. We can see that the number of reported potentially new isoforms directly depends on the error rate. Since potentially newfound isoforms were reported only if they produced completely correct alignments, we can conclude that higher error rate in Dataset 7 prevented new isoforms perfectly matching the alignments and resulting in a very small number of reported newfound isoforms. It could also be concluded that lowering the error rate to suitable levels is very important for accurately determining new isoforms in this way.

**Figure 5.**
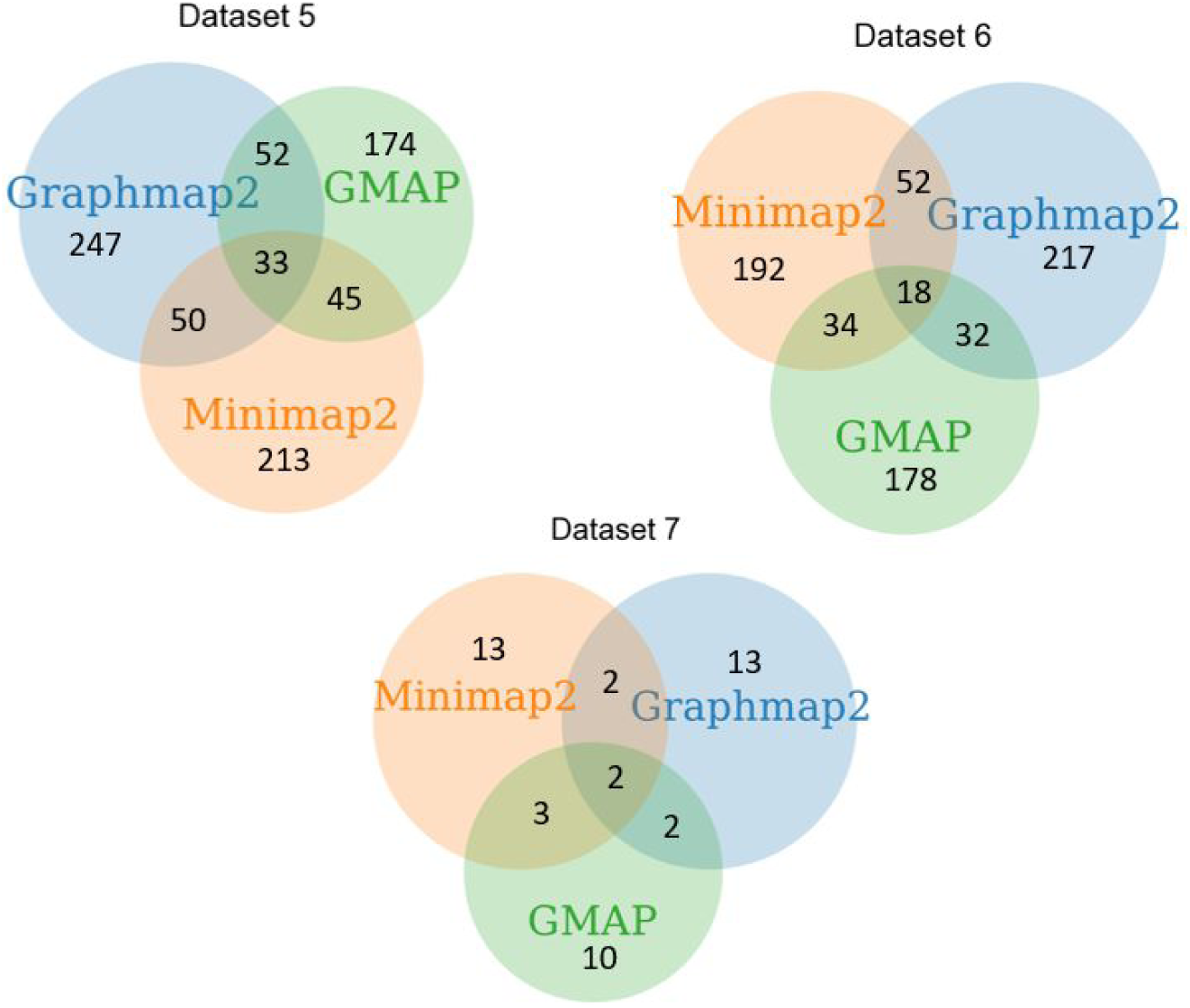
Overlapping of potentially new transcripts among mappers for each dataset.

Similar analysis can be made by looking at the results for each mapping tool. According to the results shown in Table 2, Graphmap2 maps the most reads and with the highest precision, compared to Minimap2 and GMAP. From Figure 5 we can also see that Graphmap2 produces the highest number of potentially new isoforms, with GMAP having the lowest mappability and producing the least potentially new isoforms. We can also see that a relatively small portion of determined transcripts is shared between mappers, suggesting a large number of false positives. However, Graphmap2 and Minimap2 reports slightly more transcripts shared with other mappers than GMAP. This is in accordance with the results for previously unknown gene identification.

Due to very small number of newly reported isoforms, Dataset 7 was not considered further, but for Datasets 5 and 6, overlapping of potentially new isoforms was determined for each mapper. The results are shown in Figure 6. Graphmap2 reports more newfound isoforms than Minimap2 and GMAP. Looking at datasets 5 and 6, each mapper produces a relatively large number of potential new isoforms, but only 10-20% of those were produced by all three mappers. We can conclude that analyzing new isoforms from mapping data shows interesting and promising results, but requires more work to make the results more reliable. Reported isoforms that are produced by all three mappers could be high confidence candidates suitable for further analysis.

**Figure 6.**
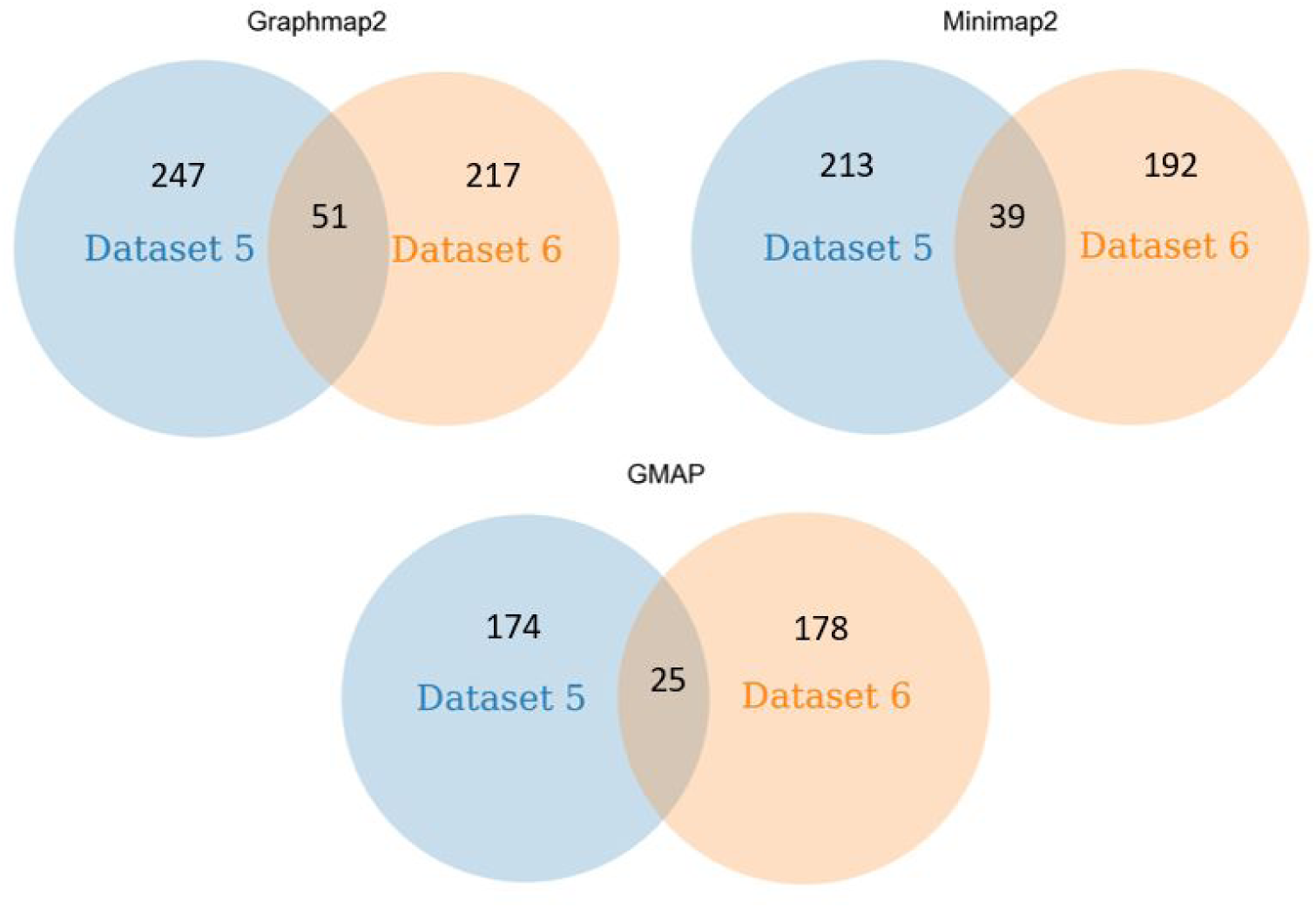
Overlapping of potentially new isoforms for each mapper.

## VI. CONCLUSION

Graphmap2 is an extended version of Graphmap, a highly sensitive tool for mapping reads to a reference genome, tailored for RNA reads. Graphmap2 creates read alignment from anchors filtered by the knapsack algorithm. It implements several alignment improvement methods, specific to RNA reads. The collective coverage information of read alignments is used to improve correctness of alignments with lower quality. Biological properties such as AG-GT donor-acceptor splice sites as well as recognition of poly(A) tail, are used to further improve the alignment.

All these improvements make Graphmap2 the very sensitive and accurate tool for RNA mapping. Evaluation on seven third-generation sequencing datasets showed that Graphmap2 has significantly higher correct measures than Minimap2 and Gmap for all datasets. Results also show that Graphmap2 is faster than Gmap but slower than Minimap2. Since Graphmap2 is based on original mapping algorithms from Graphmap optimized for earlier generation of long reads that have much higher error rate. We deem that with a less sensitive mapper we can achieve running times comparable with Minimap2 while keeping high accuracy of algorithms for the identification of exons present in transcripts and their boundaries.

Finally, we have shown that both Graphmap2 and Minimap2 have high potential in identifying previously unknown genes and isoforms. With high number of reads mapped to the same reference region by Graphmap2 and Minimap2 for which no previous annotation exists, as well as high number of donor-acceptor splice sites in alignments of these reads, Graphmap2 alignments provide indication that these alignments could belong to previously unknown genes. Similarly, results show that long reads have high potential for identification of previously unknown isoforms.

## LITERATURE

1. Jain, M., Olsen, H. E., Paten, B., & Akeson, M. (2016). The Oxford Nanopore MinION: delivery of nanopore sequencing to the genomics community. Genome Biology, 17(1), 239. https://doi.org/10.1186/s13059-016-1103-0

2. Rhoads, A., & Au, K. F. (2015). PacBio Sequencing and Its Applications. Genomics, Proteomics & Bioinformatics, 13(5), 278–289. https://doi.org/10.1016/J.GPB.2015.08.002

3. Wenger, A. M., Peluso, P., Rowell, W. J., Chang, P.-C., Hall, R. J., Concepcion, G. T., … Hunkapiller, M. W. (2019). Highly-accurate long-read sequencing improves variant detection and assembly of a human genome. BioRxiv, 519025. https://doi.org/10.1101/519025

4. Križanović, K., Echchiki, A., Roux, J., & Šikić, M. (2018). Evaluation of tools for long read RNA-seq splice-aware alignment. Bioinformatics, 34(5), 748–754. https://doi.org/10.1093/bioinformatics/btx668

5. http://bioinfo.zesoi.fer.hr/index.php/en/blog-en/56-gmap-vs-minimap2

6. Wu, T. D., & Watanabe, C. K. (2005). GMAP: a genomic mapping and alignment program for mRNA and EST sequences. Bioinformatics, 21(9), 1859–1875. https://doi.org/10.1093/bioinformatics/bti310

7. Li, H. (2018). Minimap2: pairwise alignment for nucleotide sequences. Bioinformatics, 34(18), 3094–3100. https://doi.org/10.1093/bioinformatics/bty191

8. Sović, I., Šikić, M., Wilm, A., Fenlon, S. N., Chen, S., & Nagarajan, N. (2016). Fast and sensitive mapping of nanopore sequencing reads with GraphMap. Nature Communications, 7(1), 11307. https://doi.org/10.1038/ncomms11307

9. Šošic, M., & Šikic, M. (2017). Edlib: a C/C ++ library for fast, exact sequence alignment using edit distance. Bioinformatics (Oxford, England), 33(9), 1394–1395. https://doi.org/10.1093/bioinformatics/btw753

10. Pavetić, F., Katanić, I., Matula, G., Žužić, G., & Šikić, M. (2017). Fast and simple algorithms for computing both $LCS_{k}$ and $LCS_{k+}$. Retrieved from http://arxiv.org/abs/1705.07279

11. https://github.com/lh3/ksw2

