## Supplemental materials for "Graphmap2 - splice-aware RNA-seq mapper for long reads"

### Supplementary materials

**Supplementary Table 1 Correct-5 measure results for 3 evaluated tools**

| | No. of reads | Number of aligned reads | Precision correct-5 | Recall correct-5 | F-value correct-5 |
|---|---|---|---|---|---|
| **Dataset 1** | | | | | |
| **Gmap** | 185021 | 143405 | 64.18% | 49.74% | 56.04% |
| **Minimap2** | 185021 | 174682 | 86.08% | 81.27% | 83.61% |
| **Graphmap2** | 185021 | 181507 | 85.34% | 83.72% | **84.52%** |
| **Dataset 2** | | | | | |
| **Gmap** | 411920 | 331224 | 59.21% | 47.61% | 52.78% |
| **Minimap2** | 411920 | 399449 | 61.16% | 59.31% | 60.22% |
| **Graphmap2** | 411920 | 408372 | 78.52% | 77.85% | **78.18%** |
| **Dataset 3** | | | | | |
| **Gmap** | 84007 | 63321 | 56.05% | 42.25% | 48.18% |
| **Minimap2** | 84007 | 77968 | 54.94% | 50.99% | 52.89% |
| **Graphmap2** | 84007 | 82248 | 66.03% | 64.64% | **65.33%** |
| **Dataset 4** | | | | | |
| **Gmap** | 342566 | 325080 | 28.89% | 27.42% | 28.14% |
| **Minimap2** | 342566 | 336549 | 56.56% | 55.57% | 56.06% |
| **Graphmap2** | 342566 | 341236 | 77.52% | 77.22% | **77.37%** |
| **Dataset 5** | | | | | |
| **Gmap** | 192450 | 148576 | 50.50% | 38.99% | 44.00% |
| **Minimap2** | 192450 | 181186 | 41.93% | 39.48% | 40.67% |
| **Graphmap2** | 192450 | 186885 | 54.01% | 52.45% | **53.22%** |
| **Dataset 6** | | | | | |
| **Gmap** | 243499 | 198281 | 39.23% | 31.94% | 35.21% |
| **Minimap2** | 243499 | 232116 | 42.12% | 40.15% | 41.11% |
| **Graphmap2** | 243499 | 238891 | 55.77% | 54.72% | **55.24%** |
| **Dataset 7** | | | | | |
| **Gmap** | 40318 | 37249 | 27.26% | 25.18% | 26.18% |
| **Minimap2** | 40318 | 39819 | 29.22% | 28.86% | 29.04% |
| **Graphmap2** | 40318 | 39767 | 37.55% | 37.04% | **37.29%** |

**Supplementary Table 2 Results of analysis of alignments that had not matched an annotation**

| | total number of unaligned reads | spliced reads | reads with exons next to AG-GT sites | number of neighbouring pair of exons next to AG-GT sites | number of introns |
|---|---|---|---|---|---|
| **dataset 5** | | | | | |
| **graphmap reads** | 1069 | 351 | 146 | 342 | 768 |
| **minimap reads** | 1033 | 276 | 121 | 171 | 481 |
| **gmap reads** | 929 | 457 | 104 | 208 | 2981 |
| **dataset 6** | | | | | |
| **graphmap reads** | 1014 | 313 | 150 | 272 | 558 |
| **minimap reads** | 1030 | 373 | 223 | 320 | 736 |
| **gmap reads** | 1468 | 825 | 195 | 279 | 5019 |
| **dataset 7** | | | | | |
| **graphmap reads** | 5747 | 2182 | 969 | 1075 | 2682 |
| **minimap reads** | 5732 | 1968 | 711 | 726 | 2275 |
| **gmap reads** | 5307 | 2487 | 398 | 447 | 6950 |
